# Environmental fluctuations trigger faster evolution and turnover

**DOI:** 10.1101/2025.06.04.657978

**Authors:** Pablo Duchen, Nicolas Salamin, Daniele Silvestro

**Affiliations:** Institute of Organismic and Molecular Evolution, Johannes Gutenberg University of Mainz, Hanns-Dieter-Hüsch-Weg 15, D-55128, Mainz, Germany; Institute for Quantitative and Computational Biosciences, Johannes Gutenberg University of Mainz, Johannes-von-Müller-Weg 6, D-55128, Mainz, Germany; Department of Computational Biology, University of Lausanne, CH-1015, Lausanne, Switzerland; Department of Biosystems Science and Engineering, ETH Zurich, Switzerland; Swiss Institute of Bioinformatics, 4056 Basel, Switzerland; Department of Biological and Environmental Sciences and Gothenburg Global Biodiversity Centre, University of Gothenburg, Sweden

## Abstract

Biodiversity patterns and species communities emerge from the interactions between organisms and their environment and from processes spanning across different spatial and temporal scales. Understanding the dynamics of these interactions remains difficult as it requires mapping how microevolutionary processes scale up to macroevolutionary outcomes. Here, we use an individual-based simulation framework to model how populations respond to environmental fluctuations in a spatially explicit context. We track the evolutionary processes across thousands of generations to measure the effect of emerging macroevolutionary processes under different degrees of environmental instability. We found that increased environmental fluctuations across time and space lead to elevated rates of species-level extinctions leading to homogenization of species communities, while promoting high rates of phenotypic evolution. Emerging extinction patterns indicate a selective process where species with low migration ability, narrow environmental tolerance, and limited evolvability are disproportionally affected by phases of rapid environmental change. Understanding the impact of environmental variation on organisms and species evolution remains challenging but can be explored through simulations to generate expectations and testable hypotheses about evolutionary processes across different scales.

## Introduction

The spatial distributions of species are the result of multiple ecological and evolutionary processes ultimately linked to the abiotic and biotic factors necessary for the survival of individual organisms. Species distributions are dynamic and respond to environmental changes, demographic processes, dispersal ability, and edge effects (Tomiolo and Ward 2018; Pinsky et al. 2020; Zani et al. 2024). As environments change, species must adapt to the new conditions or shift their ranges to track their environmental niches. Otherwise, they face elevated extinction risks (Tomiolo and Ward 2018). Meanwhile, the establishment of new species in recipient communities leads to species rearrangement and novel biotic interactions (Thompson and Fronhofer 2019; Calderón del Cid et al. 2024). These dynamics occur across different spatial, temporal, and taxonomic scales, encompassing population-level effects driven by short-term and localized environmental variation and species-level evolutionary adaptations and extinctions in response to long-term and large-scale changes (Donoghue and Edwards 2014).

Environmental gradients at different scales are a ubiquitous feature that shapes the species composition of biological communities. For instance, elevational gradients along mountain ranges are believed to play a crucial role in shaping biodiversity (Drummond et al. 2012; Hoorn et al. 2018; Carruthers et al. 2024) and for the survival of species during climate fluctuations, such as Quaternary Glacial Cycles (Holderegger and Thiel-Egenter 2009). Empirical studies at macroevolutionary scale have shown evidence for changes in speciation and extinction rates driven by environmental gradients and climate fluctuations (Weir and Price 2011; Silvestro and Schnitzler 2018; Condamine et al. 2013; Pino et al. 2022; Carruthers et al. 2024). Similarly, research focusing on smaller temporal and spatial scales found that these gradients and fluctuations have led to local extirpation and recolonization of populations. They have also created conditions for survival within *refugia* (Petit et al. 1997; Gavin et al. 2014), thus contributing to the biological communities that we observe today and their spatial variation. Environmental stability at low latitudes has been proposed as favorable to the accumulation of diversity (Fine 2015). Meanwhile, communities exposed to large climate fluctuations at high latitudes have been linked with higher speciation rates (Rabosky et al. 2018).

Processes of speciation, extinction, migration and evolution of traits are affected by both biotic and abiotic factors that control the distribution of populations and the fitness of individuals (Birand et al. 2011; Pimm et al. 2014; Rangel et al. 2018; Boyko and Vasconcelos 2024). Among these, the uneven distribution of species diversity has drawn attention to the complex roles of biome shifts and geographic range dynamics in driving speciation and extinction (Descombes et al. 2018; Vamosi et al. 2018). Dispersal across biomes, for instance, has been identified as a key contributor to the accumulation of biodiversity in regions such as South America (Antonelli et al. 2018). More broadly, migration and dispersal are central to many frameworks that seek to link biome transitions with evolutionary history and diversification patterns (e.g. Landis et al. 2021). In parallel, phenotypic evolution, particularly in angiosperms, has been shown to respond rapidly to environmental changes, with traits often shifting quickly following environmental novelty before stabilizing (Gorné and Díaz 2019). However, fewer studies have examined how biome shifts or long-term environmental changes influence the sustained rate of trait evolution over macroevolutionary timescales. Together, these processes shape biodiversity patterns and community composition. However, because they operate across different spatial and temporal scales, quantifying the influence of environmental gradients and their fluctuations on both micro-and macroevolutionary dynamics remains a significant challenge (Rolland et al. 2023; Calderón del Cid et al. 2024).

Simulations have proven to be powerful tools for reconstructing evolutionary processes across space and time, enabling researchers to infer patterns of biodiversity, speciation, extinction, migration, and phenotypic evolution (Hagen et al. 2021; Thompson and Fronhofer 2019; Duchen et al. 2021; Rolland et al. 2018). These approaches are particularly valuable because they allow us to “fast-forward” evolutionary dynamics that are otherwise unobservable within human timescales, and to generate expectations under explicit, mechanistic models of evolution. For example, evolutionary simulations have been used to forecast the short-term responses of coral reefs to rapid climate change over centuries (McManus et al. 2021), as well as to explore global diversification patterns over hundreds of millions of years (Leprieur et al. 2016). While macroevolutionary models are essential for identifying large-scale biodiversity patterns and estimating diversification rates, they can be limited in their ability to distinguish between underlying microevolutionary processes (Li et al. 2018). This limitation underscores the importance of integrating population-level dynamics into macroevolutionary frameworks to avoid drawing misleading causal inferences about the drivers of biodiversity.

In this study, we use individual-based simulations to assess how microevolutionary processes, operating on individuals along a fluctuating environmental gradient, scale up to macroevolutionary patterns of diversity, extinction, and trait evolution. Our simulations track individual-level traits, fitness, mortality, and dispersal, from which we derive species-level properties such as rates of migration, local and global extinction, and phenotypic evolution, so that macroevolutionary outcomes emerge from explicitly modeled mechanisms rather than being imposed as parameters. We address two research questions. First, we assess how the rapidity of the environmental change shape the spatial distribution of diversity, migration, and extinction regimes and evaluate how individual-level processes propagate the effect of the environmental shift to species-level dynamics. Second, we explore how intrinsic species properties, namely environmental tolerance, migration ability, and evolvability, modulate both, the species response to environmental change, and the rates of trait evolution and extinction at the macroevolutionary scale. By linking emergent outcomes to their underlying mechanisms, we generate theoretical expectations that can be compared against empirical data and used to formulate testable hypotheses about evolutionary processes across scales.

## Methods

### An evolutionary simulation framework

We implemented time-forward individual-based simulations in a spatially explicit context. Our simulations evolve in discrete time steps representing non-overlapping generations, in which the offspring replace the previous generation. The simulation is based on an environment divided into discrete spatial areas. The evolutionary process is based on individual-level reproduction, migration, and death. An area-specific fitness landscape affects mortality rates of the individuals based on their phenotype. The evolutionary process runs through many generations during which speciation events occur stochastically, leading to the separation of individuals into new lineages. We provide a summary of how the simulations are implemented as pseudo-code in the Supplementary Information.

### Environment

In our simulations, the environment is divided into a user-defined number *A* of discrete areas, each characterized by specific features. Each area has a carrying capacity (*K_a_*) that determines the maximum number of individuals of any species that can live in it. Furthermore, each area is characterized by a value that represents an arbitrary environmental variable (*T_a_*), such as a mean annual precipitation or temperature, for example. This value is associated with an optimal phenotype that affects the mortality of individuals (see below). The environmental variable *T_a_* can vary over time based on user-defined deterministic or stochastic trajectories. Finally, the connectivity between areas is described by a matrix *M* that controls the frequency of migration between pairs of areas. This matrix can be used to, for instance, assume equal connectivity among all areas or to allow direct migration only between adjacent ones, e.g. stepping-stone migration.

### Species

Species in our simulations are characterized by a set of features that determine their evolution in time and space. Each species has a migration parameter (*m_s_*) that describes its ability to move across areas based on their connectivity defined by the matrix *M* (see below). A growth rate (*g*) determines the mean number of offspring per individual per generation and leads to an increasing number of individuals if *g >* 1. We also include a species-specific carrying capacity (*K_s_*), which represents the maximum number of individuals that can be sustained in one area. However, the actual number of individuals of a species in an area is also determined by other factors including the fitness of its individuals and the presence of other species that contribute to the area-specific carrying capacity (*K_a_*). A species is also characterized by a fixed range of possible phenotype values, outside of which individuals have a mortality rate of 1. This property effectively limits the phenotypic space within which a species can evolve throughout its life, with parallels to the concept of phenotypic stasis (Gould and Eldredge 1977; Eldredge et al. 2005), but allowing the species mean to evolve within the range. The center of the range is set to the mean phenotype between individuals of a species at its time of origination, and the size of the range (*V*) can be set to an arbitrary number, thus modulating the potential of the species to evolve. Finally, each species is assigned a rate of phenotypic evolution (*σ_G_*) that affects the amount of expected phenotypic change between the parent and offspring.

### Evolutionary process

Our simulations evolve through discrete time steps, i.e. generations, during which four main processes take place, namely reproduction, migration, death, and speciation. At each step, each individual of generation *t* reproduces asexually and is replaced by a number of offspring sampled from a Poisson distribution Pois(*g*), where *g* is the species-specific growth rate. We model trait inheritance phenotypically, without explicitly simulating the underlying genetic architecture: the trait value of each offspring is drawn from a normal distribution centered on the trait value of its parent, with standard deviation *σ_G_*. This follows the infinitesimal model of quantitative genetics, in which offspring phenotypes are normally distributed around the parental value with a variance independent of the parental genotype (Fisher 1918; Bulmer 1980; Barton et al. 2017); under asexual reproduction, *σ_G_* thus summarizes the per-generation mutational input to phenotypic variance and sets the maximum rate at which a lineage can respond to selection (Lande 1976; Lynch and Lande 1993; Bürger and Lynch 1995). The same phenotypic approach to trait evolution is commonly adopted in spatially explicit eco-evolutionary simulation models (Rangel et al. 2018; Hagen et al. 2021; Pontarp et al. 2019).

Individuals of the new generation migrate from the starting area *i* to a new area *j* with probability *m_s_ ×* **M***_ij_*. Thus, the migration process is controlled by a species-specific dispersal capacity *m_s_* and by connectivity among areas *M*.

The probability of survival of an individual is a function of the area carrying capacity, the fitness of the individual, and its trait value relative to the phenotypic range of the species (*V*). An individual will die with probability *δ* calculated as follows:

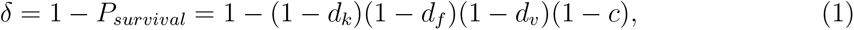

where *d_f_*and *d_k_* are the death probabilities determined by fitness and carrying capacity, respectively, *d_v_* is the mortality linked to their phenotype, and *c* is a baseline death probability (e.g. death caused by random events). For a species *s* with *n_s,a_* individuals in area *a*, let *L _s,a_* = min(*K_s_, F_a_*) denote the limiting capacity, where 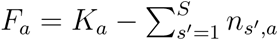 is the free space in the area based on its carrying capacity *K_a_*. The death probability due to carrying capacity is then

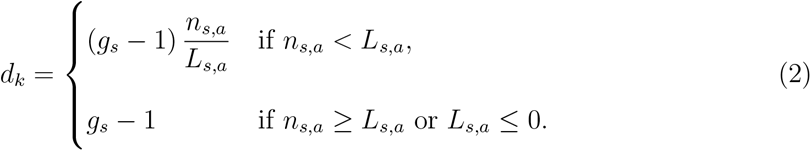

Thus, *d_k_* increases linearly with population size relative to the limiting capacity and reaches its maximum value *g_s_ −* 1 once the species reaches its species-specific carrying capacity (*K_s_*), or once the area is full, i.e. when *F_a_ ≤* 0 because of the individuals of other species. Beyond this point, *d_k_* is held at its maximum, so a population at or above its limiting capacity cannot grow in expectation. Stochastic overshoots are possible but are counteracted in subsequent generations. We used an area-specific fitness landscape to describe the fitness of individuals as a function of their phenotypic trait value (Simpson 1944; Arnold et al. 2001; Svensson 2012; Boucher et al. 2018). Specifically, the probability of death of an individual depends on the position of the phenotypic trait value in the fitness landscape, which is described by an optimum *θ_a_* and a standard deviation *σ_f_*. For simplicity, we assumed here that the optimal trait value in each area matched the environmental value *θ_a_* = *T_a_*. Thus, the fitness-based death probability of an individual with phenotype *x* in area *a* is

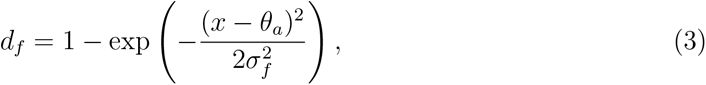

which corresponds to a Gaussian fitness function with optimum *θ_a_* and width *σ_f_*, i.e. the probability density of a normal distribution *N* (*θ_a_, σ*^2^) evaluated at *x* and rescaled to its maximum. Under this formulation, *d_f_* = 0 for an individual with optimal phenotype (i.e. *x* = *θ_a_*), and *d_f_* increases towards 1 as the phenotype departs from the optimum.

Finally, we incorporated a species-specific phenotypic tolerance component by assuming that individuals that attain trait values outside the species-specific phenotypic range die with probability *d_v_* = 1. Thus, for a species with initial mean phenotype *x*_0_ and phenotypic range 2*V*

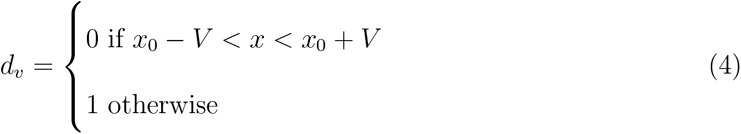

At each generation, each species will split and give rise to a new descendent species with probability

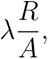

where *λ ∈* [0, 1] is the maximum probability of speciation, *R* is the number of areas occupied by the parent species, and *A* is the total number of areas. Thus, under this parameterization, we assume that speciation is more likely for geographically widespread species compared to narrowly distributed ones (Rosenzweig 2001). Speciation is assumed to occur within one area, where a random set of individuals is randomly assigned to the descendent species. As each new species originates as a population within a single area, its initial range is *R* = 1. Thus the speciation rate is assumed equal across all species at their time of origination. As species expand or shrink their range through migration and mortality, the speciation rate varies proportionally.

Each new species is randomly assigned a phenotypic range *V* around its mean phenotype (the arithmetic mean of the trait values of the initial individuals). New species are also randomly assigned a set of specific parameters, namely a growth rate (*g*), carrying capacity (*K_s_*), and a rate of phenotypic evolution (*σ_G_*). Although our simulation framework does not explicitly parameterize the extinction process, extinctions can occur if all individuals of a species die.

## Simulations under environmental shifts

### Environment parameterization

We simulated a gradient across *A* = 9 areas, with environmental values *T*_1_*, …, T*_9_ following a logistic function ranging from *T*_1_ *≈* 0 to *T*_9_ *≈* 5. Thus, for the *i^th^* area, the environmental value is

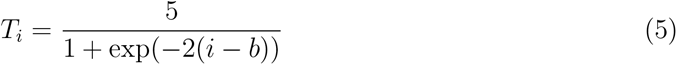

where 2 is the steepness of the function and *b* is the midpoint. Here we modeled environmental shifts by changing the midpoint *b* over time based on a bounded random walk, such that *b_t_*_+1_ *∼ N* (*b_t_, σ_E_*). We used reflection at the boundaries, set here to 2 and 8, to constrain the midpoint to occur within the simulated areas. We simulated three scenarios where environmental shifts occurred at different rates, according to *σ_E_*equal to 0.001 (slow change), 0.01 (intermediate) and 0.1 (rapid change; Fig. 1). A logistic function is a parsimonious representation of a bounded spatial environmental cline, in which environmental conditions change relatively little toward the extremes but undergo a more pronounced transition across an intermediate region. This structure is intended to approximate broad-scale ecological gradients in which distinct environmental regimes occur toward opposite ends of a spatial axis. We performed 100 simulations assuming equal connectivity among all areas (*M_i̸_*_=_*_j_* = 0.01). The area carrying capacity (*K_a_*) was fixed to 5000 individuals for all areas.

**Figure 1:**
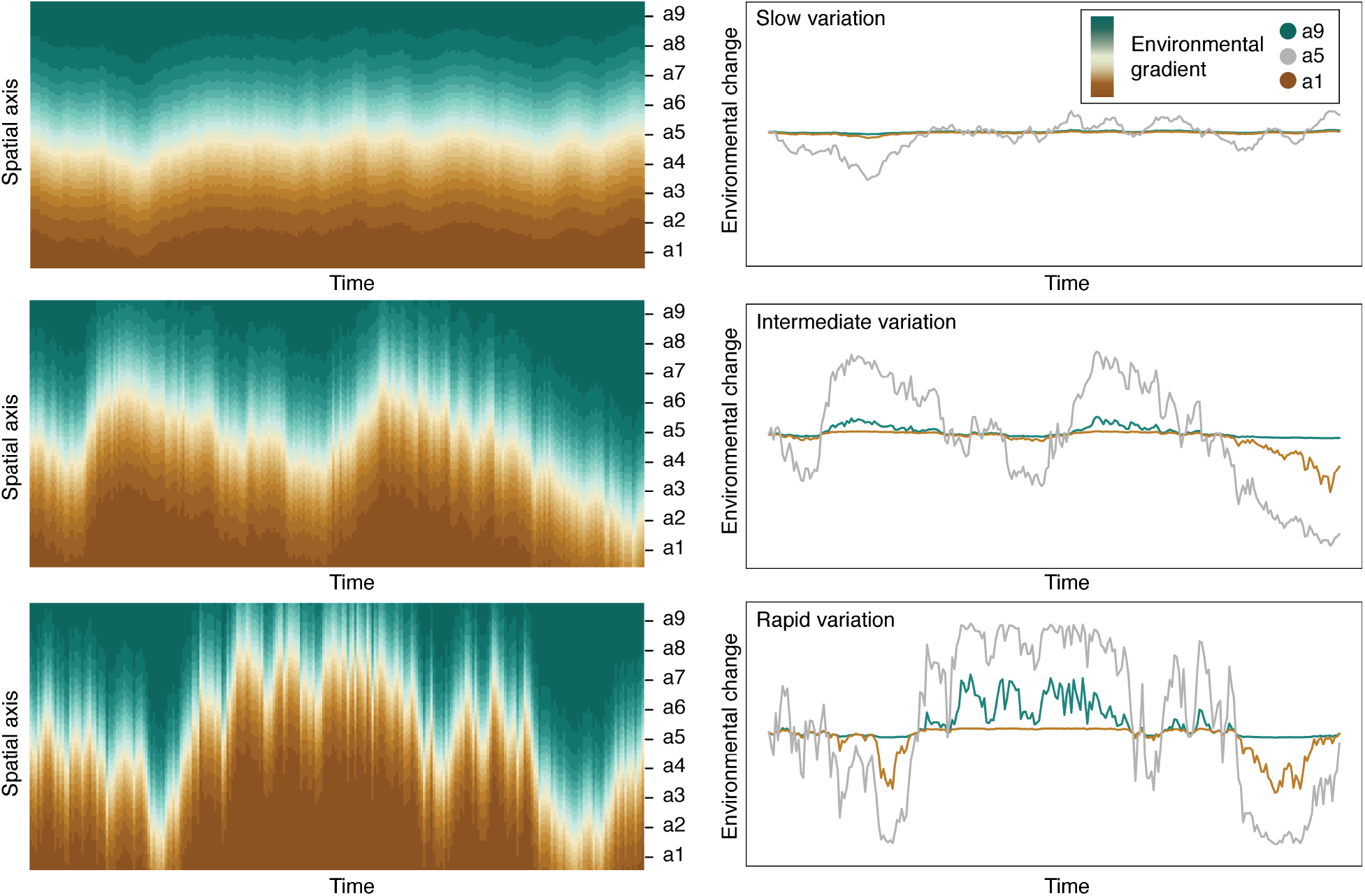
Simulated biome shifts under different rates of environmental change. The y-axis represents the nine spatial units used in our simulations that discretize an environmental gradient (areas *a*_1*−*9_). a) The environmental values in the areas varied according to a bounded random walk with different rates of change (slow, intermediate, and rapid) leading to fluctuations through time. b) The relative change in environmental value through time is shown for areas *a*_1_*, a*_5_*, a*_9_, demonstrating how the areas at the extremes are subject to lower intensity of change, while the intermediate areas fluctuate the most.

### Simulation settings

Each simulation was run for 25,000 generations starting from one species with 500 individuals occurring in one randomly selected area. Each individual was assigned a random phenotypic value drawn from a normal distribution centered on the optimal trait value for the area and a standard deviation of 0.01. We set the maximum probability of speciation *λ* to 0.0002.

For each species, including the initial one, we randomly drew species-specific parameters. The growth rates (*g*) were sampled from a uniform distribution *U* (1.1, 2). We sampled species-specific migration rates from three values set to *m_s_ ∼* {0.00001, 0.0001, 0.01}, which capture migration abilities varying by three orders of magnitude. The range of traits of the species was sampled from three possible values 2*V ∼* {1, 5, 12}, which describe species with a narrow, intermediate, and wide distribution of possible phenotypes. The rate of phenotypic evolution was randomly drawn from a uniform distribution *σ_G_ ∼ U* (0.005, 0.02). Finally, the species carrying capacity (*K_s_*) was drawn from a uniform distribution ranging between the initial population size of a species and 4,000 individuals.

### Evaluation of outcomes

We summarized the simulations to evaluate emerging patterns focusing on species richness, migration, extinction, and phenotypic evolutionary rates. We separated simulations based on the scenarios of slow, intermediate, and rapid environmental shift (*σ_E_*= 0.001, 0.01, 0.1).

We looked at the species pool at the end of each simulation and calculated diversity indices. Specifically, we computed the alpha diversity as the number of species occurring in each area. We then calculated the beta diversity as the Sørensen index averaged across all pairs of areas (Baselga and Leprieur 2015; Whittaker 1960) using the vegan R package (Dixon 2003). We also computed the gamma diversity as the number of living species at the end of the simulation. Finally, we computed the endemism richness as the number of species occurring only in one area (Kier et al. 2009).

We summarized migration patterns and computed empirical migration rates as the average number of successful migrations per species per time unit for each pair of areas. We considered migrations successful when the migrating individuals were able to survive for more than one generation, giving them the opportunity to reproduce and transmit their genetic material to the population in the destination area.

We computed empirical extinction rates as the average number of extinction events per species per time unit. These rates were calculated as local extinction rates by looking at individual areas, global extinction rates by looking at disappearance of entire species, and by separating species by the magnitude of their trait range (*V*).

We estimated the effect of environmental changes on trait evolution at the species level by inferring the rates at which species mean phenotypes change. We first computed the empirical rate of trait evolution within each area as the mean absolute difference between species phenotypes across consecutive generations. We computed species mean phenotypes as the average trait value among all individuals of the same species occurring in each area. These rates are a function of both the species-specific rates of phenotypic evolution (*σ_G_*) and the impact of fitness and the environment on survival.

Finally, we used phylogenetic comparative methods (PCM) to estimate the rate of trait evolution based on macroevolutionary models as inferred from the community of species reaching the end of each simulation. To this aim, we used the phylogenetic tree resulting from the simulation pruned to include only extant species and the species mean phenotypes at the tips of the tree as input. We used the geiger package (Pennell et al. 2014) to estimate the maximum likelihood evolutionary rates based on a Brownian motion model and phylogenetic signal using Pagel’s lambda (Pagel 1999). Estimates of evolutionary rates and phylogenetic conservatism were interpreted as descriptive summaries of the outcomes resulting from our simulations and used to assess whether the living species and their traits and phylogeny would still reveal a signature of the historical environment-driven evolutionary process.

## Results

We simulated evolutionary processes along an environmental gradient following a logistic function, where the placement of the transition between two extreme values (the midpoint) fluctuates over time. This setting could capture the transition between different adjacent environments, such as tropical forests and savannas or above and below the tree line in mountain settings.

We simulated three scenarios that differed by the rapidity with which environmental change occurs. In the slow change scenario (*σ_E_* = 0.001) the expected change per time step is on average 10 times smaller than in the intermediate change scenario (*σ_E_* = 0.01; Fig. 1) and 100 times smaller than in the rapid change scenario (*σ_E_* = 0.1).

The areas at the two extremes of the gradient are less prone to change compared to the intermediate areas where the fluctuations are stronger (Fig. 1). In the slow change scenario, most of the environmental variation is concentrated in the intermediate areas, while in the rapid change scenario all areas are affected by substantial variation through time.

We found that environmental gradients and their changes over time play a fundamental role in shaping evolutionary processes with repercussions on species diversity, longevity, and selection, and on the resulting communities. Simulations with a steep but geographically stable environmental gradient show that this acts as a permeable barrier to dispersal. We found that different degrees of environmental change and proximity to different environments have a significant impact on the processes of migration, extinction, and phenotypic evolution.

### Spatial diversity patterns

The distribution of species richness resulting from our simulations varied substantially depending on the environmental change scenario. Although speciation rates were kept constant in all simulations and equal across all areas, we found strong heterogeneities in the resulting diversity patterns. The mean gamma diversity was *∼* 77 species (s.d. 17 among simulations) in the slow change scenario (*σ_E_* = 0.001), while lower diversity was attained in the rapid change scenario (*σ_E_* = 0.1), i.e. *∼* 56 species (s.d. 14). We found that in the rapid change scenario, alpha diversity decreases towards the middle area that corresponds to where environmental fluctuations were strongest (Fig. S1). This pattern in alpha diversity was less pronounced with slower change scenarios.

Environmental change played an important role in determining the heterogeneity in species communities between areas. The median beta diversity for the slow change scenario was 0.81 (95% CI 0.77–0.84), it decreased to 0.78 (95% CI 0.72–0.82) in the intermediate scenario and to 0.72 (95% CI 0.67–0.77) in the rapid change scenario. Thus, increasing the frequency and magnitude of environmental fluctuations leads to a homogenization of species communities. A similar pattern is observed in the endemism richness that was higher and homogeneously distributed in slow change scenarios while showing substantially lower values in areas affected by strong environmental fluctuations under a rapid change scenario (Fig. 2a).

**Figure 2:**
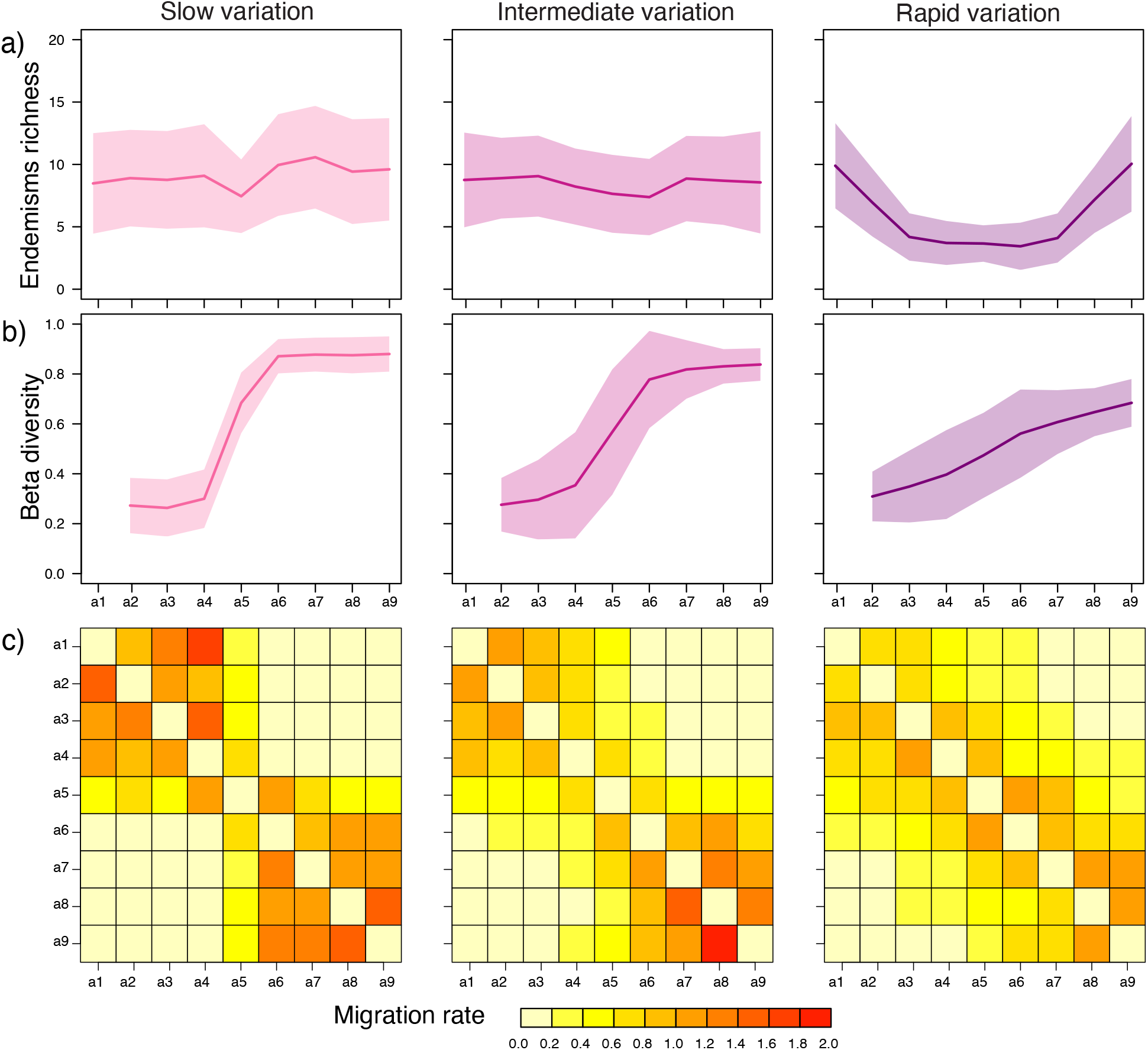
See full caption on the next page a) Distribution of endemism richness across the nine simulated areas. The number of species endemic to a single area decreases in areas with high environmental variability, e.g. *a*_2*−*8_ under rapid fluctuations, and *a*_5_ with slow fluctuations. Rapid environmental fluctuations lead to higher levels of homogenization of the species composition with generally fewer endemics. b) Beta diversity (Sørensen index) under different regimes of environmental variation (from slow to rapid) calculated between the area at one environmental extreme (*a*_1_) against all others along the gradient. Although the magnitude of the environmental gradient is equal across all scenarios, beta diversity increases at a lower slope under rapid fluctuations. This indicates that rapid environmental fluctuations lead to a homogenization of spatial biodiversity patterns. c) Migration rates were computed as the mean number of successful migrations per species per 1000 generations among pairs of areas, where we considered as successful any migration event that resulted in the species surviving in the area for more than one generation. In the presence of slow environmental fluctuations, successful migration rates reach the highest level especially within environmentally similar areas e.g. *a*_1*−*4_ and *a*_6*−*9_ while these are more rare across steeper environmental gradients e.g. from *a*_3_ to *a*_6_. As the environmental fluctuations become more rapid, the distribution of migrations comes to resemble a stepping-stone scenario, primarily connecting environmentally adjacent areas, even though there were no geographic distances implemented in these experiments and the *a priori* probability of dispersal was set equal among all pairs of areas.

**Figure 3:**
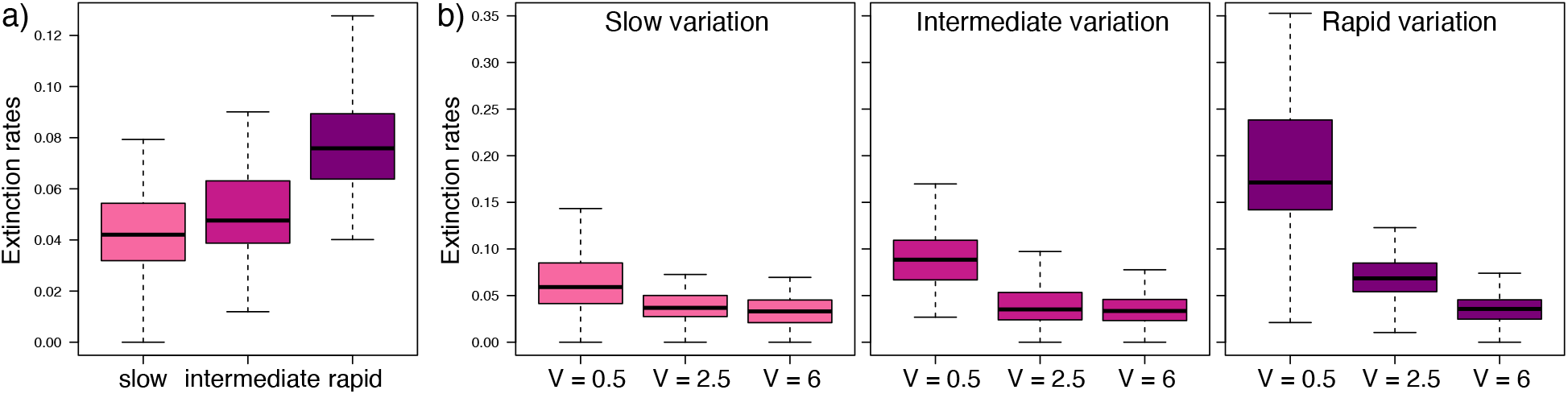
Extinction rates (expected number of extinction events per species per 1000 generations) under different regimes of environmental variation (from slow to rapid). a) Overall extinction rates are substantially higher under rapid environmental fluctuations compared with slow fluctuations. b) This pattern is associated with an elevated extinction rate for species with narrow tolerance range (here 2*V* = 1, 5), while species with wider tolerance (2*V* = 12) remain largely unaffected.

### Migration patterns

The rates of migration are a function of species migration ability, area connectivity and ability of the individuals of a species to successfully establish in a new area (i.e. to survive in the area for more than one generation). Our simulations showed that environmental fluctuations did not affect the overall rates of migration, with negligible difference between rapid and slow change scenarios. Yet, the spatial distribution of migrations differed strongly among scenarios. In simulations with slow change migrations were concentrated within regions with similar environmental values, e.g., among areas 1–4 and among areas 6–9, based on the simulated logistic gradient. This reflects the fact that individuals adapted to the environment in, e.g., area 1 have a low chance to survive and establish in an area with different environmental value (e.g. area 6), even though the underlying connectivity and dispersal ability would allow them to migrate there. Thus, the environmental gradient creates a permeable barrier. In contrast, in rapid change scenarios, migrations are less spatially concentrated but distributed across areas (Fig. 2c), reflecting the rapid fluctuations in the placement of the permeable barrier. These spatial patterns of migration contribute to explaining the decreasing beta diversity (i.e. increasing homogenization of species communities) observed with increasing rapidity of environmental change.

### Patterns and magnitude of extinction

Even though our model did not include a species-level extinction rate, we observed extinction events across all simulations. These are related to individual mortality due to fitness and carrying capacity of the areas. Extinction rates were overall 60% higher under rapid environmental change compared to intermediate change scenarios and 82% higher compared to slow change scenarios.

When disaggregating extinction rates based on the magnitude of the species’ tolerance range (i.e. variable *V*), we found that species with wide tolerance (2*V* = 12) had the lowest extinction rates and their rates were similar regardless of the environmental change scenarios. In contrast, species with a small tolerance range (2*V* = 1) were more prone to extinction and negatively affected by the rapidity of environmental change. For instance, in slow change scenario species with small-tolerance range showed an extinction rate 80% higher than wide-tolerance species, whereas in rapid change scenarios the increase was almost 5-fold.

Extinction rates were also strongly correlated with the ability of the species to migrate. In particular, we found that species with high migration ability showed the lowest extinction rates with little variation related to the rapidity of environmental change. In species with high dispersal capacity, the mean extinction rate was 0.037 (species per 1000 generations) under strong fluctuations and 0.030 with more stable environments (Fig. S3). In contrast, slow-migrating species had a mean extinction rate of 0.159 under strong fluctuations (4-fold increase) and 0.060 under small fluctuations (2-fold increase).

Extinction rates are always higher in scenarios with rapid environmental change compared to slow change. However, in species with low evolvability (here *σ_G_ ∈* [0.004, 0.006]), the average extinction rates increased by 47.6% in simulations with rapid environmental change compared to those with slow change. In contrast, in fast-evolving species (here *σ_G_ ∈* [0.018, 0.020]) the average extinction rates increased only by 16.8%. Thus, slow-evolving species are disproportionately affected by rapid environmental change relative to fast-evolving ones.

As extinction disproportionally affected species with narrow tolerance and low dispersal ability, rapid environmental changes led to favor species with a wider geographic range (mean range across species: 3.11 areas) compared with slow environmental change (mean range across species: 2.48 areas). This, in turn, affected the realized speciation rates (given that the probability of speciation was modeled as proportional to range size), which therefore were on average higher in simulations with rapid environmental changes (Fig. S4a). These higher rates compensate for the elevated extinction, as shown by the nearly identical net diversification rates and total simulated number of species across all scenarios (Figs. S4c, S5b). Yet, after rescaling the realized speciation rates by the average range size across each environmental change scenario, we recover essentially equal speciation rates, reflecting the assumption in our simulations of constant speciation probability per area per species (Fig. S4b).

### Phenotypic evolution

In our simulations, the phenotype was represented by a trait that evolved randomly from the parent to the offspring. As the value of the trait determined the fitness and mortality rate of the individual (depending on the local environmental variable), the mean phenotype of a species evolved by tracking its optimum based on the geographic distribution of the species. The empirical rates of phenotypic evolution calculated within each area showed that, at the macroevolutionary scale, environmental variation leaves a clear signature on traits. Empirical rates were up to one order of magnitude higher in areas with a high magnitude of environmental change compared with more stable areas (Fig. S2).

Changes in phenotypic rate of evolution varied across scenarios also at the species level, i.e. taking the average phenotype for each species. The median rate of phenotypic evolution calculated based on phylogenetic comparative methods under a Brownian model of evolution was 1.6 times higher under rapid environmental change compared to intermediate change and 2.5 times higher compared to slow change scenarios (Fig. 4a).

**Figure 4:**
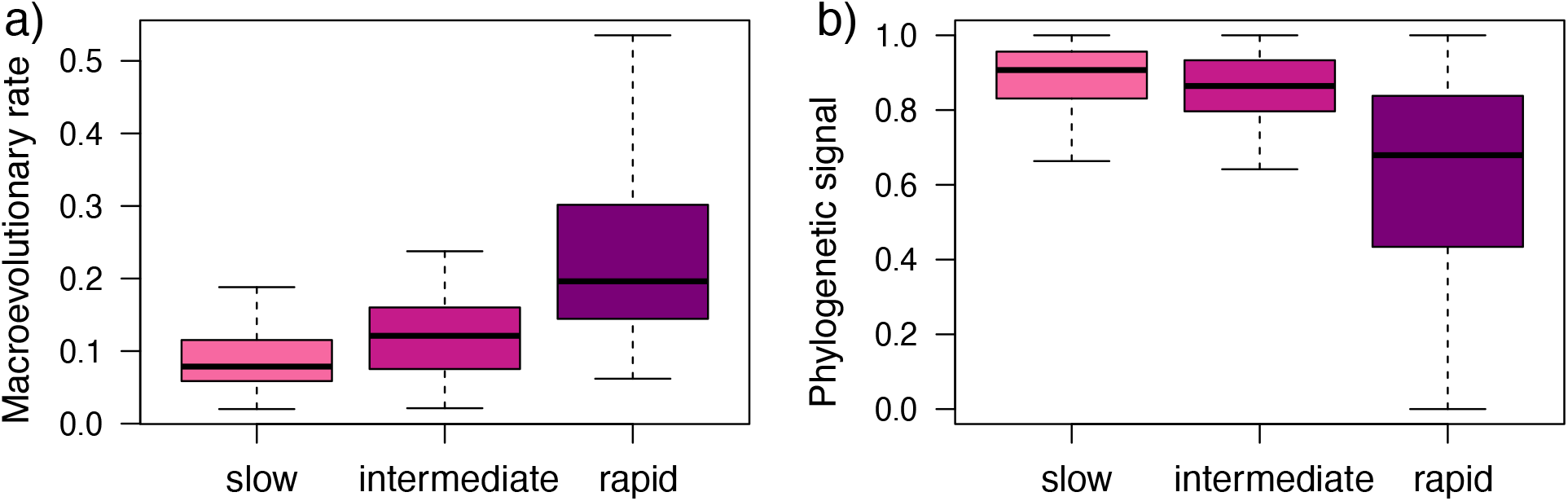
Estimated macroevolutionary parameters using phylogenetic comparative methods. a) The evolutionary rates assuming a Brownian model show that increasing rapidity of environmental variation leads to substantially faster phenotypic evolution at the species level. b) In contrast, the degree of phylogenetic signal, here measured using the lambda statistic (Pagel 1994), strongly decreases with rapid environmental change.

Environmental variation also had the effect of generally reducing the levels of phylogenetic signal in the trait data, as quantified by the lambda parameter. This generally exceeded 0.8 in stable environments, indicating phylogenetically conserved traits, while it ranged between 0.4 and 0.8 in rapidly changing scenarios, with the distribution of phenotypes becoming more constrained by changes in fitness over time than by phylogenetic inheritance.

## Discussion

We presented a simulation framework tracking microevolutionary processes at the individual level and scaling up to macroevolutionary scales, leading to speciation and extinction of entire lineages (Vrba and Eldredge 1984). Our simulations show that environmental fluctuations, an extrinsic forcing acting only on individual survival, propagate to the species level as elevated extinction, homogenized communities, and accelerated realized rates of trait evolution, and that intrinsic species properties (tolerance range, migration ability, and evolvability) determine which lineages bear the cost of these fluctuations.

Our simulations include a representation of an ecological gradient characterized by a single variable and of geography based on a set of discrete areas. Although there are more complex and realistic simulation methodologies (Hagen et al. 2021), these typically focus on broader biogeographic and evolutionary processes and longer-term climate change that unfolds over many millions of years and is shaped by plate tectonics and the configuration of land masses and ocean basins (Skeels et al. 2023a). These approaches have led to significant insights into the build-up of global biodiversity patterns such as the latitudinal diversity gradient and turnover across major biogeographic areas (Skeels et al. 2023b). The scale of these simulations means that some individual-level features, such as fitness and migration, are approximated to a population level within grid cells and that small-scale environmental variation through time and space might be averaged out. Alternatively, simulations designed to capture fine-scale processes provide useful tools to e.g. model biodiversity responses to climate change in relatively short amounts of time (McManus et al. 2021), but might not be able to scale up and capture macroevolutionary processes. In contrast, we focused here on finer spatial and temporal scales while still being able to derive macroevolutionary and macroecological patterns emerging from our individual-based simulations. The simulated environmental gradient and its fluctuations over time can be interpreted as an abstract representation of common boundaries between adjacent ecosystems, such as the primarily temperature-driven tree line along elevation gradients (Büntgen et al. 2022) or the precipitation-driven transitions between forest and open habitats (Ivory et al. 2018; Andermann et al. 2022). This framework can therefore capture processes occurring at relatively short timescales, such as glacial cycles or regional changes of precipitation patterns. These processes directly impact the fitness, survival, and dispersal ability of the individuals, with consequences scaling up to shaping the fate of entire species.

### Consequences of environmental fluctuations on migration and extinction patterns

Our simulations predict that environmental fluctuations leave clear signatures in the spatial distribution of species diversity and have a strong impact on migration patterns. For example, the level of endemism is comparatively high and roughly uniformly distributed when biodiversity evolves in stable environments, while it is strongly reduced in areas affected by a high magnitude and frequency of environmental variation. This supports empirical observations and theoretical expectations linking environmental stability with the ability of ecosystems to accumulate higher diversity (e.g. Ackerly 2003; Wiens et al. 2010; Crisp and Cook 2012). This concept has been used to explain broad-scale patterns such as the latitudinal diversity gradient (Wiens and Donoghue 2004), with, e.g., stable tropical forests characterized by extremely high diversity, including staggering numbers of rare species (ter Steege et al. 2013) in contrast to species-poor northern European forests that have been intensely and repeatedly affected by Quaternary glacial cycles (Svenning 2003).

Simulations based on low degrees of environmental change over time led to 40% higher diversity at present compared to simulations performed under rapid environmental fluctuations (Fig. S5a). However, our simulations used the same speciation probability (proportional to geographic range size) across species, and the resulting total number of species including living and extinct is similar across all simulations (Fig. S5b). Thus, the lower present diversity associated with high environmental variation appears to be primarily driven by increased extinction, which was substantially higher under strong environmental variation and disproportionately affecting species with small ranges linked to narrow tolerance and low migration ability. Because the probability of speciation in our model depends not only on the fixed maximum speciation parameter *λ* but also on a species’ occupied range (*R/A*), differences in geographic range among simulation scenarios also influence the realized speciation rates (Jablonski 1986; Payne and Finnegan 2007). In our simulations, extinction selectivity favoring species with wider occupied range led to higher realized speciation rate that (Fig. S4). This essentially led to simulations with higher species turnover, that explains the similarity in the total number of species produced across scenarios.

In our framework, we did not explicitly parameterize extinction with a predefined rate, and the disappearance of species emerged from environmentally driven mortality of individuals in the simulation. Thus, extinction emerged from the ecological and evolutionary mechanisms incorporated into the model rather than from a predefined extinction rate. Rapid environmental change can increase the mismatch between the phenotype of individuals and the local environmental optimum, thereby increasing fitness-related mortality (*d_f_*). In addition, individuals whose phenotypes fall outside the species-specific phenotypic range experience complete mortality (*d_v_* = 1). Increased individual mortality can consequently reduce population sizes sufficiently for species to disappear. Thus, the higher extinction rates observed under rapid environmental fluctuations emerge from the interaction between environmental change, phenotypic adaptation, and the demographic constraints imposed by carrying capacity and species-specific phenotypic ranges. Our findings are consistent with empirical evidence linking lower diversity and higher extinction rates in regions that have historically been affected by strong climatic fluctuations, as observed, for example, in the tree diversity of central and northern Europe, likely driven by Quaternary glaciation cycles (Svenning and Skov 2007; Zhang et al. 2022; Gamboa et al. 2024). These predictions are also in line with the patterns of increased biodiversity turnover observed in time series over the past few decades of climate change and anthropogenic changes to the environment (Pinsky et al. 2025).

The patterns described here are, to some extent, dependent on the time scale considered and on the level of connectivity among areas. In our simulations, we chose to allow for dispersal with equal probability across areas. However, in other contexts, environmental fluctuations can lead to the emergence and removal of barriers, such as changes in sea level, microclimate *refugia*, or drainage rearrangements in fluvial systems, all of which can spur dispersal and species diversification (Albert et al. 2018; Salles et al. 2021; Morales-Barbero et al. 2018). Thus, the net effect of environmental oscillations on migration and species diversification is likely context-dependent, and simulations with micro- to macroevolutionary dynamics like those presented here can help us make predictions under different scenarios.

### Linking trait evolution and environmental change across scales

One of the emerging properties of our simulations is that phenotypic macroevolutionary rates, as commonly inferred through phylogenetic comparative methods at the species level (Harmon 2019), are affected by environmental fluctuations.

In our analyses, we found that the estimated macroevolutionary rates increased with stronger environmental fluctuations, while the phylogenetic signal decreased. These expectations for adaptive traits may be used to create testable hypotheses in empirical datasets based on phylogenetic comparative methods. For instance, likelihood-based methods can be used to compare the macroevolutionary rates and phylogenetic conservatism (Revell et al. 2018) between clades hypothesized to have experienced stable or variable environments. The use of simulations and statistical inference can thus provide us with independent tests for eco-evolutionary hypotheses.

Our simulations assumed that phenotypic variation within a species is generated through a random process with a constant species-specific variance between the parent and the offspring, which reflects the intrinsic evolvability of the species. However, when measuring the rate of phenotypic change at macroevolutionary scales, we found that it increased substantially with increasing environmental variation over time (Figs. 4, S2).

This indicates that macroevolutionary rates are the combined result of evolvability on the microevolutionary scale and environmental change (Tsuboi et al. 2024; Latrille et al. 2024; Duchen et al. 2020). Discrepancies between the micro-and macroevolutionary dynamics are supported by ample empirical evidence (Uyeda et al. 2011; Rolland et al. 2018; Scholl and Wiens 2016; Henao Diaz et al. 2019) and fit theoretical expectations (e.g. Rolland et al. 2023; Duchen et al. 2021). Our simulations coupling microevolutionary changes with a dynamic environment show that the macroevolutionary outcomes are indeed a function of both the intrinsic evolvability of a species and external forces.

An important consideration is that our model assumes a constant variance in trait inheritance (*σ_G_*) within species. Therefore, the accelerated rates of phenotypic evolution observed under environmental instability should not be interpreted as changes in the underlying rate at which phenotypic variation is generated. Rather, these rates emerge from the interaction between stochastic phenotypic variation and environmental selection. As environmental optima fluctuate, changes in the fitness landscape can alter the phenotypic composition of surviving populations and consequently produce larger changes in species mean phenotypes across generations. The macroevolutionary rates estimated from the simulated phylogenies should therefore be interpreted as realized rates of phenotypic evolution under changing environmental conditions. Exploring models in which the variance of trait inheritance itself varies among species or in response to environmental conditions, as predicted by individual-based models with explicit simulation of genetic changes (Romero-Mujalli et al. 2021), would be a useful extension of the framework.

### Life-history traits and extinction risks

The prime driver of extinction in our simulations was the environmentally driven variation in the fitness of individuals that resulted in changes in their mortality rate, in some cases leading to the demise of a whole species (Vrba 1993). Three main factors emerged as having a major impact on determining the eventual extinction of a species, namely the breadth of a species environmental tolerance, its migration ability, and the species evolvability. Species with a narrow tolerance range or low ability to move or adapt were the most prone to extinction, in particular in rapidly changing environments. The relationship between tolerance breadth and extinction risk is also consistent with the resource-use hypothesis, which links ecological specialization to differences in speciation and extinction rates (Vrba 1987; Hernández Fernández and Vrba 2005; Cantalapiedra et al. 2011). These emerging properties are in line with theoretical expectations and empirical evidence from both modern and past biodiversity dynamics (Forester et al. 2022; Jablonski 2022; Harte et al. 2004; Hollenbeck and Sax 2024; Araújo et al. 2005; Peniston et al. 2020), with potential implications for conservation strategies (Gillson et al. 2013; Pinsky et al. 2022, 2025). As biodiversity is facing the most intense, rapid, and widespread environmental change probably ever experienced by any of the living species today, tolerance range, ability to adapt or track suitable habitat will likely prove crucial to determining species survival (Wiens and Zelinka 2024; Pinsky et al. 2025).

### Model limitations and future development

A simplification of our framework concerns the speciation process. While extinction emerges from individual-level ecological and evolutionary processes, speciation is implemented using a common baseline parameter *λ*, with its probability additionally scaled by geographic range. The framework could readily be extended by allowing *λ* to vary among species as a function of species characteristics such as migration capacity, phenotypic variability, or demographic properties. More mechanistically, speciation probability could also depend on ecological heterogeneity or phenotypic differentiation among populations occupying different areas. Such extensions would allow speciation, like extinction, to respond more directly to ecological and evolutionary conditions emerging during the simulation. We retained a common *λ* in the present study to maintain a parsimonious model and isolate the processes investigated here.

Our framework also involves several simplifying assumptions regarding the spatial and ecological processes represented. Area-specific fitness landscapes provide a simple representation of spatially varying selection, whereby different environmental conditions favor different phenotypes. However, real fitness landscapes are likely to depend on multiple abiotic and biotic factors and may not be well described by a single optimum. In our modeling of migration, the simulation does not explicitly represent the time or energetic costs associated with physical movement, nor does it distinguish among different dispersal distances or account for detailed establishment processes following colonization. A stepping-stone structure (or any other migration structure represented by *M*) should therefore be viewed as a simplified representation of spatial connectivity and individual dispersal. These assumptions make the framework tractable for exploring the interaction between environmental change, migration, and evolutionary dynamics, but also mean that predictions should be interpreted at the spatial and temporal scale represented by the model.

Our results can also be viewed in the context of rate-mismatch theory, which proposes that extinction occurs when environmental change exceeds the capacity of species to adapt. Recent theoretical work has represented this capacity as an effective species-level adaptation rate integrating genetic, phenotypic, ecological, demographic, and geographic processes (Rothman and Petrovskii 2026). In our framework, this capacity is not specified as a single parameter but emerges from the interaction among phenotypic variability (*σ_G_*), tolerance range (*V*), migration capacity (*m_s_*), geographic range, demographic constraints, and spatial structure. Thus, our individual-based model provides a mechanistic perspective on how species-specific tracking capacities may arise and how they can determine persistence under changing environments. A useful extension would be to quantify an emergent tracking capacity for each simulated species and examine its distribution and relationship to extinction risk. Because our simulations are not calibrated to an absolute time scale, such a comparison would be most appropriately formulated in terms of dimensionless environmental change per generation relative to species-level tracking capacity, rather than absolute rates.

## Conclusions

Understanding how spatial and temporal environmental variation affects individual organisms, populations, and in the long run the fate of entire species is a challenging yet fundamental aim of evolutionary research. Our knowledge of the mechanisms at the interface between environment and organisms at different temporal and spatial scales can benefit from theoretical work (Rosindell et al. 2010; Rosenzweig and McCord 1991; Calderón del Cid et al. 2024) and advances in the analysis of empirical data under increasingly realistic models (Ezard et al. 2011; Quintero and Landis 2020; Gaboriau et al. 2025). Modeling the relationships between environmental changes and micro- and macroevolution, rates of phenotypic change, extinction and species accumulation in space and time however, remains difficult (Rolland et al. 2023). Our study shows that these links can be explored through simulations, leading to emerging patterns that can be used to formulate testable hypotheses and improve our interpretation of the evolutionary processes at different scales.

## Code and data availability

All code was written in R. Code available at: https://github.com/phylolab/phylotraitsim

## Supplementary Information

### Supplementary text

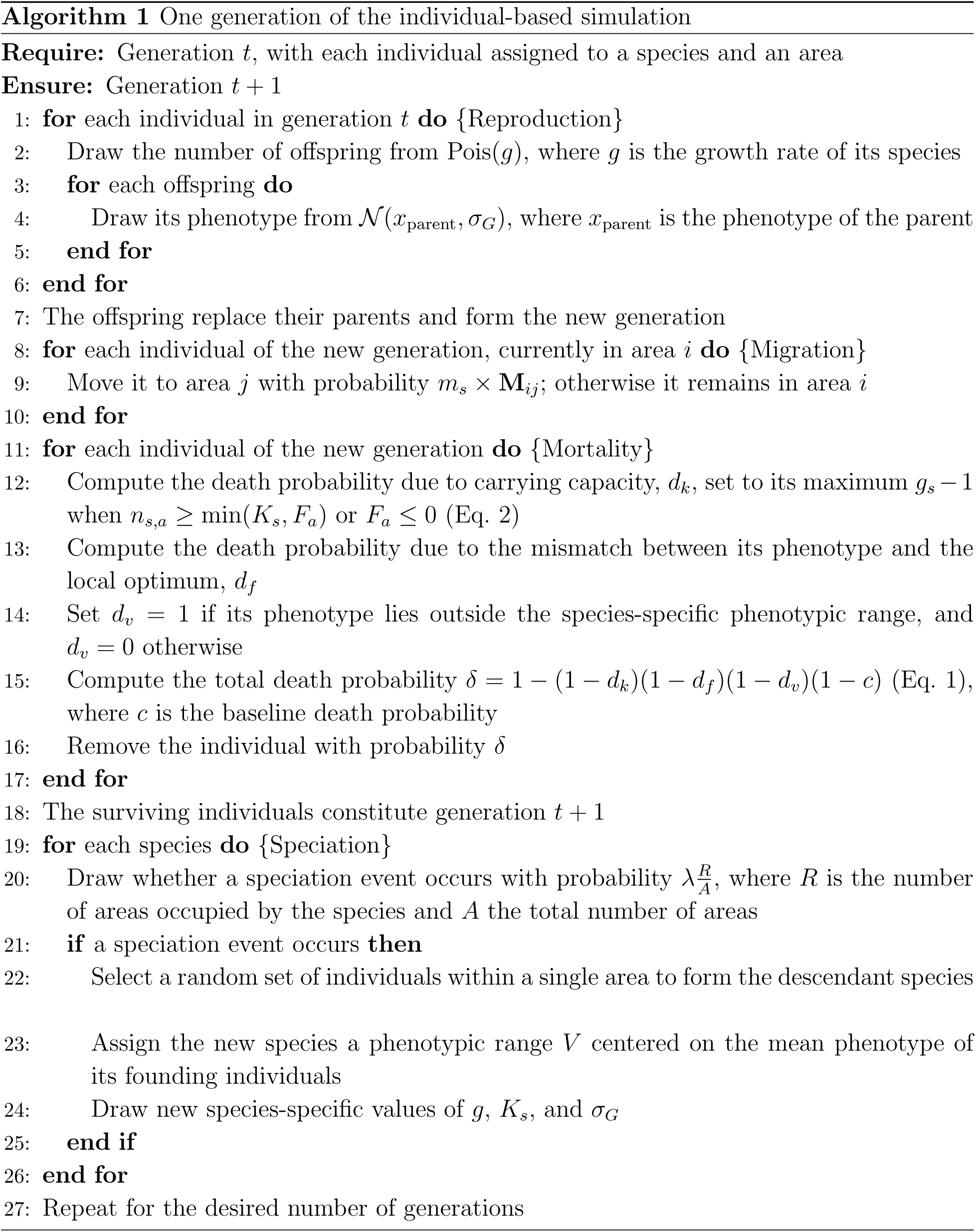

|  |  |
|---|---|
| **Require:** | Generation *t*, with each individual assigned to a species and an area |
| **Ensure:** | Generation *t* + 1 |
| 1: | **for** each individual in generation *t* **do** {Reproduction} |
| 2: | Draw the number of offspring from Pois(*g*), where *g* is the growth rate of its species |
| 3: | **for** each offspring **do** |
| 4: | Draw its phenotype from 𝒩(*x*<sub>parent</sub>, σ<sub>*G*</sub>), where *x*<sub>parent</sub> is the phenotype of the parent |
| 5: | **end for** |
| 6: | **end for** |
| 7: | The offspring replace their parents and form the new generation |
| 8: | **for** each individual of the new generation, currently in area *i* **do** {Migration} |
| 9: | Move it to area *j* with probability *m*<sub>*s*</sub> × **M**<sub>*ij*</sub>; otherwise it remains in area *i* |
| 10: | **end for** |
| 11: | **for** each individual of the new generation **do** {Mortality} |
| 12: | Compute the death probability due to carrying capacity, *d*<sub>*k*</sub>, set to its maximum *g*<sub>*s*</sub> − 1 when *n*<sub>*s*,*a*</sub> ≥ min(*K*<sub>*s*</sub>, *F*<sub>*a*</sub>) or *F*<sub>*a*</sub> ≤ 0 (Eq. 2) |
| 13: | Compute the death probability due to the mismatch between its phenotype and the local optimum, *d*<sub>*f*</sub> |
| 14: | Set *d*<sub>*v*</sub> = 1 if its phenotype lies outside the species-specific phenotypic range, and *d*<sub>*v*</sub> = 0 otherwise |
| 15: | Compute the total death probability δ = 1 − (1 − *d*<sub>*k*</sub>)(1 − *d*<sub>*f*</sub>)(1 − *d*<sub>*v*</sub>)(1 − *c*) (Eq. 1), where *c* is the baseline death probability |
| 16: | Remove the individual with probability δ |
| 17: | **end for** |
| 18: | The surviving individuals constitute generation *t* + 1 |
| 19: | **for** each species **do** {Speciation} |
| 20: | Draw whether a speciation event occurs with probability $\lambda\frac{R}{A}$, where *R* is the number of areas occupied by the species and *A* the total number of areas |
| 21: | **if** a speciation event occurs **then** |
| 22: | Select a random set of individuals within a single area to form the descendant species |
| 23: | Assign the new species a phenotypic range *V* centered on the mean phenotype of its founding individuals |
| 24: | Draw new species-specific values of *g*, *K*<sub>*s*</sub>, and σ<sub>*G*</sub> |
| 25: | **end if** |
| 26: | **end for** |
| 27: | Repeat for the desired number of generations |

### Supplementary figures

**Figure S1:**
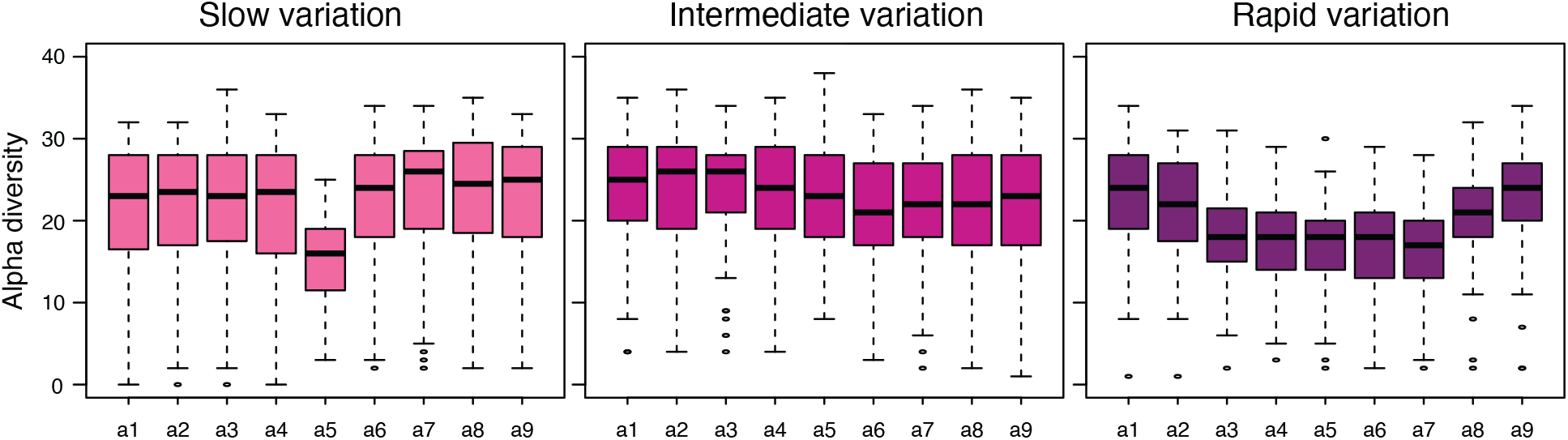
Number of living species in each area at the end of the simulation.). Areas subjected to higher frequency and magnitude of change (*a*_5_) are linked with lower alpha diversity, with the effect being more strongly pronounced in simulations with rapid environ-mental variation

**Figure S2:**
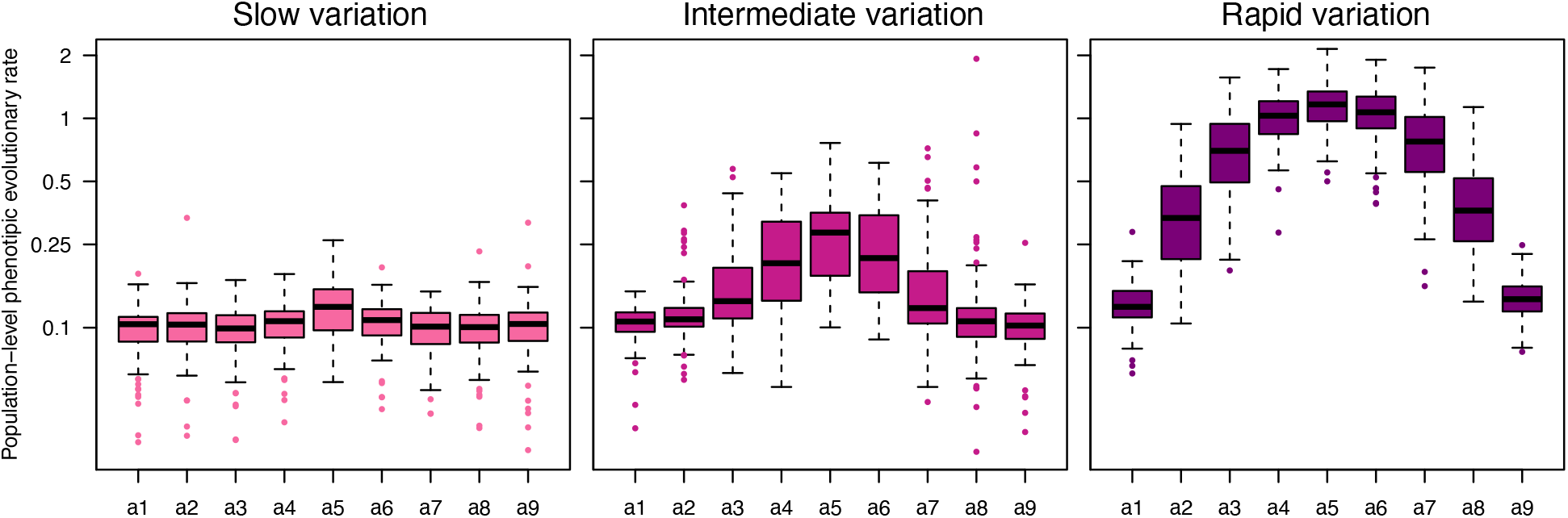
Empirical rates of phenotypic trait evolution computed within each area across different scenarios of environmental variation (from slow to rapid). The rates are computed separately for each area, as the absolute variation in phenotypic mean at the population level, i.e. including all individuals of each species, across generations. Here the rate of change is reported using 1000 generations as a time unit and was log10 transformed. Areas subjected to higher frequency and magnitude of change (*a*_5_) are linked with more rapid trait evolution relative to other areas, as the result of average phenotypes being driven by fitness. As observed based on phylogenetic comparative methods Fig. 4, rates of phenotypic evolution are overall substantially higher in simulations with strong environmental variation.

**Figure S3:**
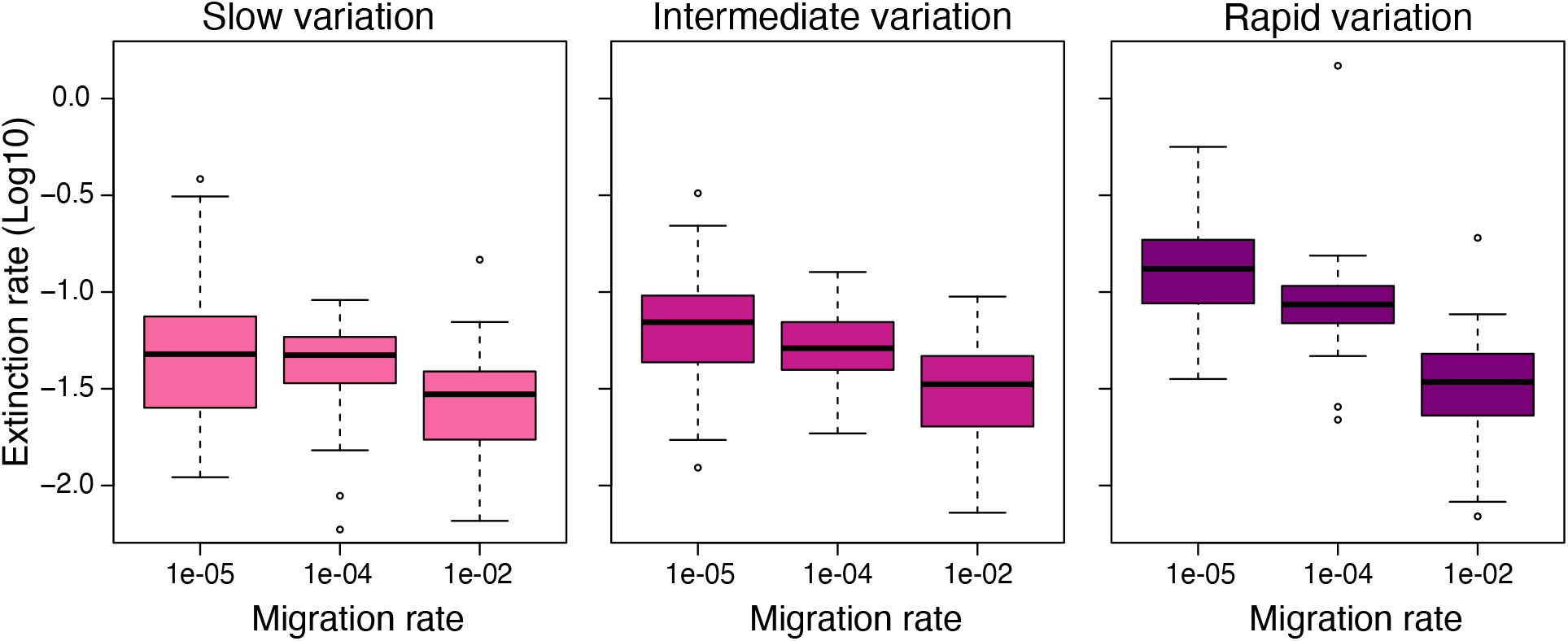
Extinction rates (log10 transformed) plotted against species migration ability under different regimes of environmental variation (from slow to rapid). Ability to migrate allows species to track their environmental optima thus reducing their extinction probability.

**Figure S4:**
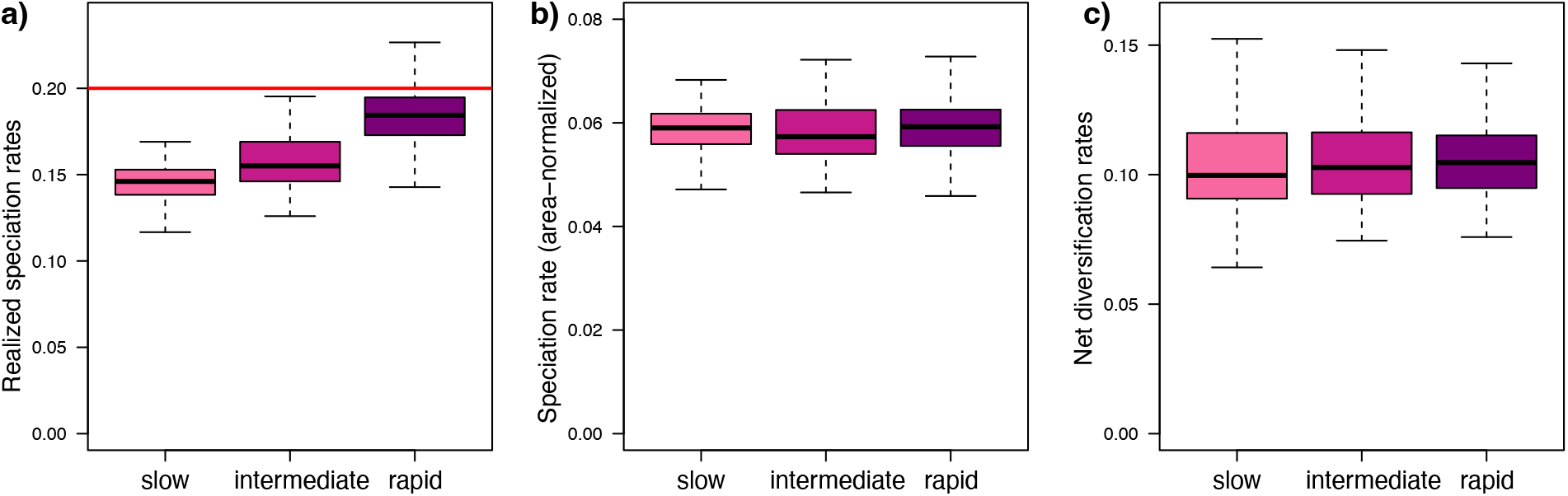
Realized speciation, extinction, and net diversification rates under different regimes of environmental variation (from slow to rapid). The rates use 1000 generations as time unit. Rapid environmental change led to disproportional extinction of species with small geographic range, which in turn affects the species probability to speciate up to a maximum rate here set to 0.2 (red line). Thus, the realized speciation rates were higher rapid environmental change, with increases matching the higher extinction rates leading to roughly equal net diversification rates.

**Figure S5:**
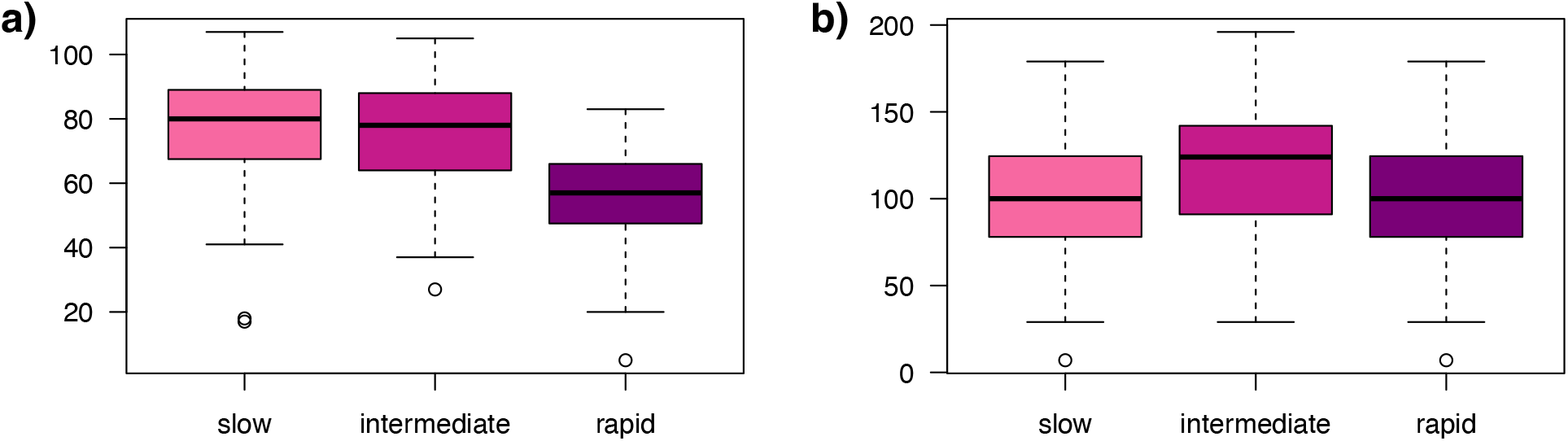
a) Total number of living species at the end of the simulations run under different regimes of environmental variation (from slow to rapid). b) Total number of species living and extinct. These were found to be consistent among simulations with different environmental variation, reflecting corresponding diversification rates shown in Fig. S4.

### Supplementary tables

**Table S1:** Parameters and variables of the evolutionary simulation model, including their meaning, values or sampling distributions, units/scales, and potential calibration to empirical data. Parameters are grouped into fixed simulation settings, species-specific parameters drawn at the origination of each species, and derived or state variables computed during the simulation.

| **Symbol** | **Meaning** | **Value / range / sampling distribution** | **Unit / scale** | **Potential calibration to real data** |
|---|---|---|---|---|
| *Fixed simulation settings* |  |  |  |  |
| *A* | Number of discrete spatial areas | *A* = 9 | Number of areas; discrete | Set according to the number of empirical spatial units. |
| *K*<sub>*a*</sub> | Area-specific carrying capacity | *K*<sub>*a*</sub> = 5000 | Individuals per area | Estimate from population-size or density data. |
| *M*<sub>*ij*</sub> | Connectivity between areas *i* and *j* | *M*<sub>*i*≠*j*</sub> = 0.01 (equal connectivity) | Dimensionless connectivity matrix | Estimate from geographic distance, landscape resistance, or movement data. |
| *T*<sub>*a*</sub> | Environmental value of area *a*; determines the local phenotypic optimum | $T_a = \frac{5}{1+e^{-2(a-b_t)}}$, ranging from *T*<sub>1</sub> ≈ 0 to *T*<sub>9</sub> ≈ 5; the steepness of the logistic function is fixed to 2 | Environmental-variable units | Use measured environmental data and rescale to the model range. |
| *b*<sub>*t*</sub> | Midpoint of the environmental gradient at generation *t* | *b*<sub>*t*+1</sub> ∼ 𝒩(*b*<sub>*t*</sub>, *σ*<sub>*E*</sub>), reflected at the boundaries 2 and 8; initial value *b*<sub>0</sub> = … | Spatial-area units | Estimate from the spatial position of the environmental gradient and fit its temporal trajectory. |
| *σ*<sub>*E*</sub> | Magnitude of environmental change per generation | {0.001, 0.01, 0.1} (slow, intermediate, and rapid change, respectively) | Environmental-gradient units per generation | Estimate from temporal environmental variation. |
| *λ* | Maximum probability of speciation per generation | *λ* = 0.0002 | Probability per generation | Estimate from phylogenetic diversification rates. |
| *c* | Baseline probability of death | *c* = 0.00001 | Probability per generation | Estimate from baseline mortality/survival data. |
| *t* | Generation (time step) | 0, …, 25000 | Generations; discrete | Relate one generation to the empirical generation time. |
| – | Initial population | One species with 500 individuals in one randomly selected area; phenotypes drawn from 𝒩(*θ*<sub>*a*</sub>, 0.01) | Individuals; phenotypic trait units | Initialize from observed abundance and trait distributions. |
| – | Number of replicate simulations | 100 per environmental-change scenario | Simulations | — |
| *Species-specific parameters (drawn at the origination of each species)* |  |  |  |  |
| *g*<sub>*s*</sub> | Growth rate; mean number of offspring per individual per generation | 𝒰(1.1, 2) | Offspring per individual per generation | Estimate from demographic or recruitment data. |
| *K*<sub>*s*</sub> | Species-specific carrying capacity | 𝒰(*n*<sub>*s*,0</sub>, 4000), where *n*<sub>*s*,0</sub> is the population size of the species at origination | Individuals per species per area | Estimate from species abundance time series. |
| *m*<sub>*s*</sub> | Migration capacity | {0.00001, 0.0001, 0.01} | Dimensionless probability multiplier | Estimate from observed movement or migration frequencies. |
| 2*V* | Width of the species-specific phenotypic range, centered on *x*<sub>0</sub> | {1, 5, 12} (narrow, intermediate, and wide range, respectively) | Phenotypic trait units | Estimate from observed physiological or ecological tolerance ranges. |
| *σ*<sub>*G*</sub> | Rate of phenotypic evolution; standard deviation of the offspring phenotype around the parental value | 𝒰(0.005, 0.02) | Phenotypic trait units per generation | Estimate from parent–offspring trait variation. |
| *σ*<sub>*f*</sub> | Standard deviation of the fitness landscape around the optimum | 𝒰(10, 100) | Phenotypic trait units | Estimate from the empirical phenotype–fitness relationship. |
| *x*<sub>0</sub> | Mean phenotype of a species at origination; centre of its phenotypic range | Arithmetic mean of the trait values of the founding individuals | Phenotypic trait units | Calculate from observed trait values of founding individuals. |
| *Derived quantities and state variables* |  |  |  |  |
| *x* | Individual phenotypic value | Continuous; offspring phenotype drawn from 𝒩(*x*<sub>parent</sub>, *σ*<sub>*G*</sub>) | Phenotypic trait units | Use observed individual trait measurements. |
| *θ*<sub>*a*</sub> | Optimal phenotype in area *a* | *θ*<sub>*a*</sub> = *T*<sub>*a*</sub> | Phenotypic trait units | Estimate from empirical phenotype–environment relationships. |
| *n*<sub>*s*,*a*</sub> | Number of individuals of species *s* in area *a* | Emergent variable | Individuals per species per area | Use abundance data to calibrate and validate the model. |
| *F*<sub>*a*</sub> | Free space available in area *a* | *F*<sub>*a*</sub> = *K*<sub>*a*</sub> − ∑<sub>*s*</sub> *n*<sub>*s*,*a*</sub> | Individuals per area | Calculate from empirical carrying capacity and abundance. |
| *R* | Number of areas occupied by a species | 1, …, *A* | Number of areas; discrete | Estimate from species occur-rence/occupancy data. |
| *i*, *j* | Source and destination area indices | 1, …, *A* | Discrete spatial index | Correspond to empirical spatial units. |
| *d*<sub>*k*</sub> | Death probability due to carrying-capacity limitation | $\frac{(g_s-1)\,n_{s,a}}{\min(K_s,F_a)}$ for *n*<sub>*s*,*a*</sub> < min(*K*<sub>*s*</sub>, *F*<sub>*a*</sub>) (Eq. 2) | Probability per generation | Calibrate jointly with *K*<sub>*a*</sub>, *K*<sub>*s*</sub>, and *g*<sub>*s*</sub> using abundance data. |
| *d*<sub>*f*</sub> | Death probability due to phenotypic mismatch with the local optimum | $1-\frac{f(x\|\theta_a,\sigma_f)}{f(\theta_a\|\theta_a,\sigma_f)}$ | Probability per generation | Estimate from survival or fitness as a function of phenotype. |
| *d*<sub>*v*</sub> | Death probability outside the species-specific phenotypic range | 0 if *x*<sub>0</sub>−*V* < *x* < *x*<sub>0</sub>+*V*; 1 otherwise | Probability per generation | Estimate from empirical trait tolerance limits. |
| *δ* | Total probability of death | 1 − (1 − *d*<sub>*k*</sub>)(1 − *d*<sub>*f*</sub>)(1 − *d*<sub>*v*</sub>)(1 − *c*) (Eq. 1) | Probability per generation | Fit to observed individual survival/mortality. |

